# Loss of T73T75 phosphorylation on PP2A-B56^Par1^ advances mitotic entry and reduces *S. pombe* cell size

**DOI:** 10.64898/2026.01.19.700315

**Authors:** Lenka Hálová, Yvonne Connolly, Duncan L Smith, Janni Petersen, Iain M Hagan

## Abstract

Mitosis is triggered when the rising activity of CDK1-Cyclin B, amplified by the CDK1/Cdc25/Wee1 feedback loop, overcomes inhibitory signalling from Wee1 and counteracting phosphatases. CDK-opposing phosphatases PP1, PP2A-B55 and PP2A-B56 are regulators of mitosis. A screen for differentially phosphorylated sites in a *ΔPP1^dis2^* genetic background in *Schizosaccharomyces pombe* identified phosphorylation of T73 or T75 in the regulatory B56^Par1^ subunit. The B56^Par1^.T73T75 phosphorylation is directly mediated by CDK1-Cyclin B, and a phospho-mimetic mutation increased PP2A-B56^Par1^ phosphatase activity. Blocking B56^Par1^.T73T75 phosphorylation reduced cell length in unperturbed divisions from 14 to 12 µm, with no other detectable phenotypes. Therefore, blocking phosphorylation at T73T75 alone prematurely unlocked amplification of the CDK1/Cdc25/Wee1 feedback loop, advancing cells into mitosis. Signalling from T73T75 reveals for the first time that timely mitotic commitment in unperturbed cycles is mediated by PP2A-B56.

## Introduction

During interphase, CDK1-Cyclin B activity is restrained through inhibitory phosphorylation of CDK1 by Wee1 family kinases, until Cdc25 phosphatases remove the inhibitory phosphate to enforce mitotic entry (Nurse, 1990; Stern & Nurse, 1996). Cdk1-Cyclin B associates with the fission yeast equivalent of the centrosome, the spindle pole body (SPB), from early G2 until Cyclin B destruction in mitosis (Alfa *et al*, 1990; Basu *et al*, 2020; Decottignies *et al*, 2001). CDK1-Cyclin B engages Polo kinase on the SPB in a positive CDK1/Cdc25/Wee1 feedback loop 30 min before mitosis, activating the SPB (Grallert *et al*, 2013a; Grallert *et al*, 2013b; Izumi & Maller, 1993, 1995; Mulvihill *et al*, 1999). Mitosis in the whole cell is triggered when the signal amplification exceeds the inhibitory impact of Wee1 and CDK-counteracting phosphatases (Ferrell, 2008; Gould & Nurse, 1989; Hagan & Grallert, 2013; Novak & Tyson, 1993).

The Serine/Threonine phosphatases PP1 and PP2A are important regulators of mitosis in *Metazoa* and fission yeast (Nasa & Kettenbach, 2018; Nilsson, 2019; Zeisner *et al*, 2025). PP1 monomers achieve specificity by docking to regulatory proteins, which direct PP1 to substrates (Bollen *et al*, 2010; Moura & Conde, 2019). PP2A holoenzymes combine a catalytic and a scaffold subunit with one of four B-types regulatory subunits, which determine the phosphatase specificity. Of these, the B55 and B56 families have been linked to mitotic control (Lacroix *et al*, 2022; Nilsson, 2019). All three phosphatases employ preferences for the phosphorylated residue and the surrounding amino acids and are recruited to targets through specific sequences that include short linear motifs (SLiMS; (Cundell *et al*, 2016; Fowle *et al*, 2021; Hertz *et al*, 2016; Hoermann *et al*, 2020; Kruse *et al*, 2020; Padi *et al*, 2025; Smith *et al*, 2019; Zeisner *et al*., 2025)). Bulk activities of PP1 and PP2A are inhibited during mitosis, although local activities to control phosphorylations remain (Grallert *et al*, 2015; Nasa & Kettenbach, 2018; Nijenhuis *et al*, 2014; Nilsson, 2019).

In human cells, PP2A complexity is expanded by four genes encoding B55 subunits (α, β, γ, δ) and five genes encoding B56 subunits (α, β, γ, δ, ε), which utilize distinct structural folds to dictate substrate specificity (Cho & Xu, 2007; Vallardi *et al*, 2019; Xu *et al*, 2008). Furthermore, alternative splicing - particularly within the B56 genes - further expands this repertoire, enabling the formation of over 100 unique heterotrimeric combinations (McCright *et al*, 1996; Sandal *et al*, 2021). The PP2A-B55 family is the main antagonist of CDK phosphorylations and contributes to the bistability of mitotic entry and exit (Kamenz *et al*, 2021; Mochida *et al*, 2009; Nasa *et al*, 2020). PP2A-B56 and PP1 are primarily employed in local controls during mitosis, where they support microtubule-kinetochore capture, chromosome alignment and spindle assembly checkpoint silencing (Kitajima *et al*, 2006; Kruse *et al*, 2013; Smith *et al*., 2019).

In contrast to animal cells, *S. pombe* only has one B55 isoform, *B55^pab1^*, and two B56 isoforms, *B56^par1^*and *B56^par2^*, of which *par1* is the major isoform (Jiang & Hallberg, 2000). This relative simplicity makes *S. pombe*, an established eukaryotic cell cycle model, ideal for investigating conserved PP2A functions in mitosis. As in animal cells, PP2A-B55^Pab1^ dephosphorylates most CDK substrates phosphorylations (Zeisner *et al*., 2025), and couples cell growth to the cell cycle via the Greatwall-Endosulfine switch that inactivates PP2A-B55^Pab1^ at the onset of mitosis (Chica *et al*, 2016). The major PP1^Dis2^ isoform restricts the CDK1/Cdc25/Wee1 feedback loop activation on the SPB in G2 (Basu *et al*, 2022; Grallert *et al*., 2013a). During mitosis, PP1^Dis2^ and to a lesser extent PP2A-B56^Par1^ contribute to spindle assembly checkpoint silencing (Deng *et al*, 2022). All three phosphatases join forces at mitotic exit to orchestrate orderly substrates dephosphorylations (Grallert *et al*., 2015). As Cyclin B degrades, PP1^Dis2^ auto-catalytically recovers and drives a phosphatase relay, in which PP1^Dis2^ binds to and sequentially activates PP2A-B55^Pab1^ and PP2A-B56^Par1^. Binding assays and sequence conservation suggest that PP1 regulates PP2A-B55 and PP2A-B56 activities in a variety of signalling contexts throughout Eukaryotes (Grallert *et al*., 2015; Heidari *et al*, 2023). A recent systematic *in vivo* analysis of phosphorylations targeted by PP1, PP2A-B55, PP2A-B56 and the dual-specificity phosphatase CDC14 in *S. pombe* substantiated that all these phosphatases negatively regulated CDK1-Cyclin B activity (Zeisner *et al*., 2025). Interestingly, each phosphatase targeted distinct subgroups of substrate sites that were net phosphorylated at different points during *S. pombe* cell cycle, suggesting that these phosphorylation thresholds were important for mitotic onset.

The regulatory interplay between PP1^Dis2^, PP2A-B55^Pab1^ and PP2A-B56^Par1^ at mitotic exit indicates that PP1^Dis2^ targets a phosphorylation site on the respective PP2A holoenzymes (Grallert *et al*., 2015). As the regulatory subunit determines PP2A specificity, we asked whether the PP1^Dis2^ target on PP2A-B56^Par1^ resides within the regulatory subunit itself. Here, a screen for phosphorylation sites altered upon the removal of PP1^Dis2^ in fission yeast identified T73T75 phosphorylation on B56^Par1^. Mimicking T73T75 phosphorylation increased the phosphatase activity of the intact PP2A-B56^Par1^ complex, while blocking phosphorylation restricted activity. Phospho-blocking mutations altered mitotic entry, leading to premature mitosis at a reduced cell size. The T73T75 regulation thus represents a mechanism of mitotic entry control by PP2A-B56.

## Results

### PP2A-B56^Par1^.T73T75 is phosphorylated and under PP1^Dis2^ control *in vivo*

To map phosphorylation on the regulatory B56^Par1^ subunit, we first optimized large scale precipitation of B56^Par1^.HA under denaturing conditions, a method that preserves phosphorylations (Appendix Fig. S1A (Grallert & Hagan, 2017a)). To identify PP1^Dis2^-responsive sites, we then applied these optimized conditions to precipitate B56^Par1^.HA from *S. pombe* cells with or without genetic disruption of *PP1^dis2^*(Appendix Fig. S1B) and compared phosphorylation levels between samples by Quantitative Mass Spectrometry. This analysis identified differential phosphorylations within the unstructured N-terminal arm (Threonines 73 or 75 and Serine 99) and in the core B56 domain (Serine 358 or Serine 359 or Threonine 361) of B56^Par1^ (Fig. 1A – B and Supplementary Data 1). As phosphorylation of T73 or T75 was upregulated in cells lacking *PP1^dis2^*, these sites were selected for further investigation: both residues are flanked by Prolines, suggesting a cell cycle-dependent regulation (Songyang *et al*, 1994), and the high level of sequence homology surrounding these sites among B56 orthologs in the *Schizosaccharomyces* species (Fig. 1C) as well as across other Eukaryotes (Fig. 1D), supports the existence of a conserved regulatory site.

**Figure 1.**
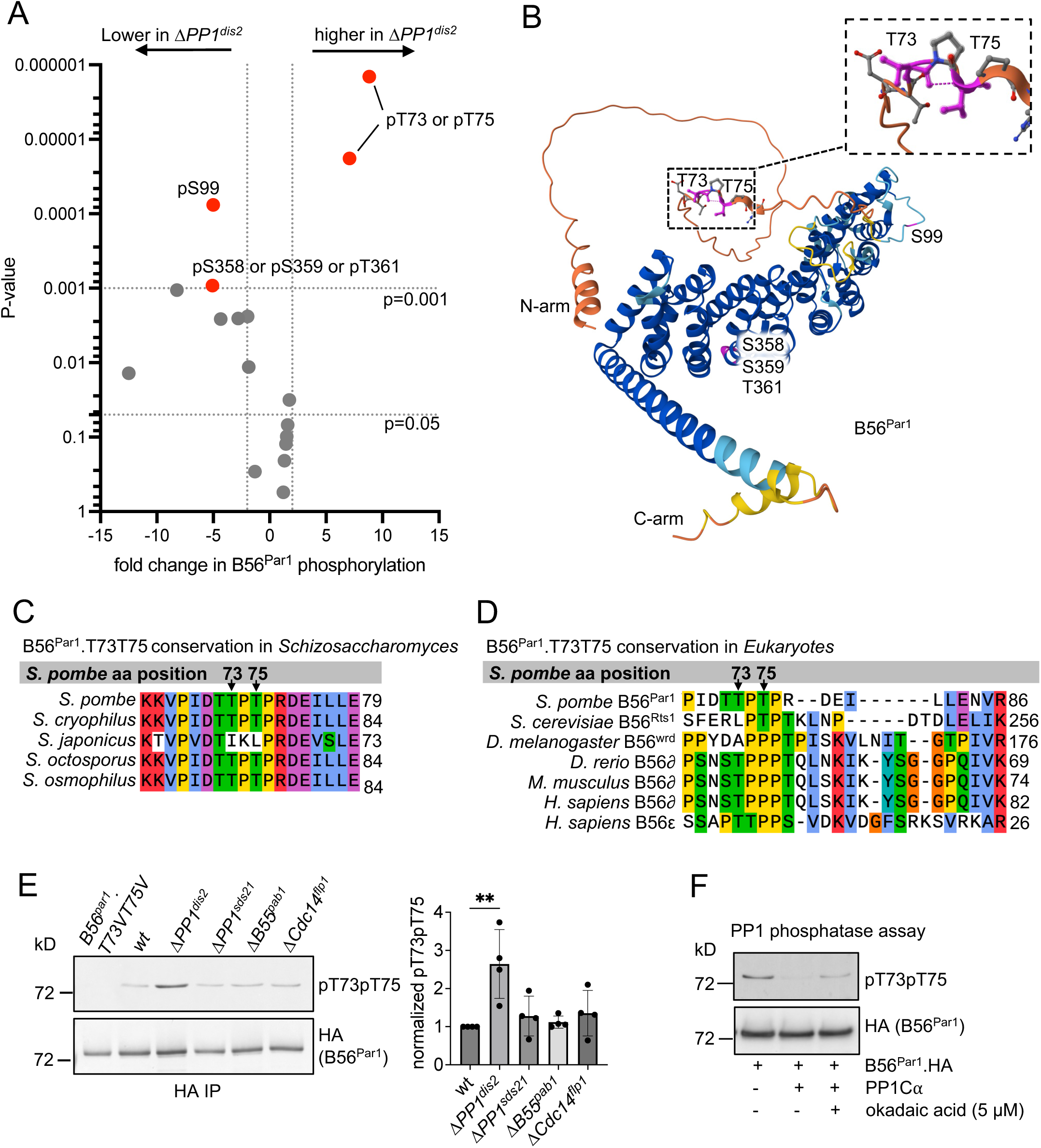
PP2A-B56^Par1^.T73T75 phosphorylation is under PP1^is2^ control. (**A**) Quantitative Mass Spectrometry Post-Translational Modifications (PTM) analysis of the impact of *PP1^dis2^* ablation upon phosphorylation of B56^Par1^. Phospho-sites changing >2fold between *PP1^dis2+^* and *ΔPP1^dis2^*(*p*<0.001) are labeled in red. (**B**) B56^Par1^ predicted structure (source: https://pombase.org) with phospho-sites identified in (**A**) highlighted in magenta. Note that T73T75 (inset) resides in a structurally disordered N-terminal arm of B56^Par1^. (**C-D**) Clustal Omega alignment of N-termini of B56 orthologs from *Schizosaccharomyces* (**C)** and *Eukaryotes* (**D**) species. (**E**) B56^Par1^.T73T75 phosphorylation levels in phosphatase-deficient mutants. Left, a representative immuno-blot; right, quantification of phosphorylation levels (means +/- standard deviation [SD], *n*=4). Phospho-specific signals detected with pT73pT75 phospho-specific antibodies were normalized to total levels of the B56^Par1^ target protein run on a parallel Western blot as sample processing controls, and normalized values were plotted relative to wild-type, which was set to 1. Significance was calculated by a two-way ANOVA followed by Dunnett’s multiple comparisons test. **P < 0.01. wt, wild type. (**F**) Dephosphorylation of B56^Par1^.T73T75 by PP1 in an *in vitro* phosphatase assay. pT73pT75 levels were detected by phospho-specific antibodies and signal was developed by a chromogenic method, and the same membrane was then probed with anti-HA antibodies as a loading control with signal developed by chemiluminescence. Okadaic acid was used as a PP1 inhibitor.

To validate the Quantitative Mass Spectrometry screen results, we generated dual phospho-specific antibodies against B56^Par1^.T73T75 and monitored phosphorylation levels in HA-immunoprecipitates from *B56^par1^.HA* cells following ablation of individual mitotic phosphatases. Importantly, the phosphorylation was elevated in cultures which lacked *PP1^dis2^*, but not other mitotic phosphatases (Fig. 1E).

Combined, these results identified a new phosphorylation site on the B56^Par1^ subunit, T73T75, which is under the regulation of the PP1 phosphatase, specifically the PP1^Dis2^ isoform.

### PP1^Dis2^ directly dephosphorylates PP2A-B56^Par1^.T73T75 *in vitro*

We next asked if B56^Par1^.T73T75 was a direct PP1^Dis2^ target. To assess whether PP1 can directly dephosphorylate T73T75, we purified B56^Par1^.HA from *S. pombe* under denaturing condition by HA affinity and used the purified protein as a substrate in phosphatase assays with recombinant PP1Cα. PP1Cα directly dephosphorylated B56^Par1^.T73T75, while the reaction was inhibited by the addition of the PP1 inhibitor okadaic acid (Fig. 1F). This result established that PP1 can directly target B56^Par1^.T73T75 *in vitro*.

We have previously shown that, at mitotic exit and as part of the PP1-PP2A phosphatase relay (Grallert *et al*., 2015), PP1^Dis2^ directly binds B56^Par1^ via G/SILK/R and K/RxVxF motifs located within the conserved common B56 core, and that mutating both motifs (*B56^par1^.PDSN*, Appendix Figure S1C) abolished this interaction (Grallert *et al*., 2015). To determine whether PP1^Dis2^-mediated dephosphorylation of B56^Par1^.T73T75 depends on PP1^Dis2^ docking to B56^Par1^, we monitored the level of T73T75 phosphorylation in the *B56^par1^.PDSN* mutant. We reasoned that if the dephosphorylation requires PP1^Dis2^ docking to B56^Par1^, the docking site mutant would mimic the elevated T73T75 phosphorylation seen in *ΔPP1^dis2^* (see Fig. 1E). Interestingly, the phosphorylation of B56^Par1^.PDSN remained at wild-type levels (Appendix Fig. S1D), indicating that in cells, PP1^Dis2^ does not target B56^Par1^.T73T75 phosphorylation via docking to the G/SILK/R and K/RxVxF motifs in B56^Par1^.

These results show that although PP1^Dis2^ can directly dephosphorylate B56^Par1^.T73T75 *in vitro*, the dephosphorylation in cells is independent of the PP1^Dis2^ docking site on B56^Par1^. Therefore, T73T75 is unlikely to represent a direct PP1^Dis2^ target within the PP1-PP2A mitotic phosphatase relay (Grallert *et al*., 2015), but instead defines a novel PP1^Dis2^-dependent mode of PP2A-B56^Par1^ regulation.

Both Threonines T73 and T75 are flanked by Prolines (see Fig. 1C-D). Since TP sites are often targeted by CDKs to regulate cell cycle progression (Songyang *et al*., 1994), we next asked if the phosphorylation is cell-cycle regulated.

### PP2A-B56^Par1^.T73T75 phosphorylation rises in G2 and peaks in mitosis

To monitor the phosphorylation in an unperturbed cell cycle, we followed cells as they progressed through cell division in cultures synchronized by centrifugal elutriation. *S. pombe* cells grown in standard minimal media divide after already completing S phase and are born in G2 phase of the cell cycle (Mitchison & Nurse, 1985). Centrifugal elutriation isolates these small G2 cells, which can then be followed over two subsequent cell division cycles, with the first cycle still showing stress response, while the second cycle represents an unperturbed division (Hagan *et al*, 2016a). B56^Par1^.T73T75 phosphorylation levels rose as septation (monitored with calcofluor staining) of the first division cell division cycle began to cease and then peaked just before the second round of division, indicating that T73T75 phosphorylation rises in G2 and is maximal during mitosis (Fig. 2A).

**Figure 2.**
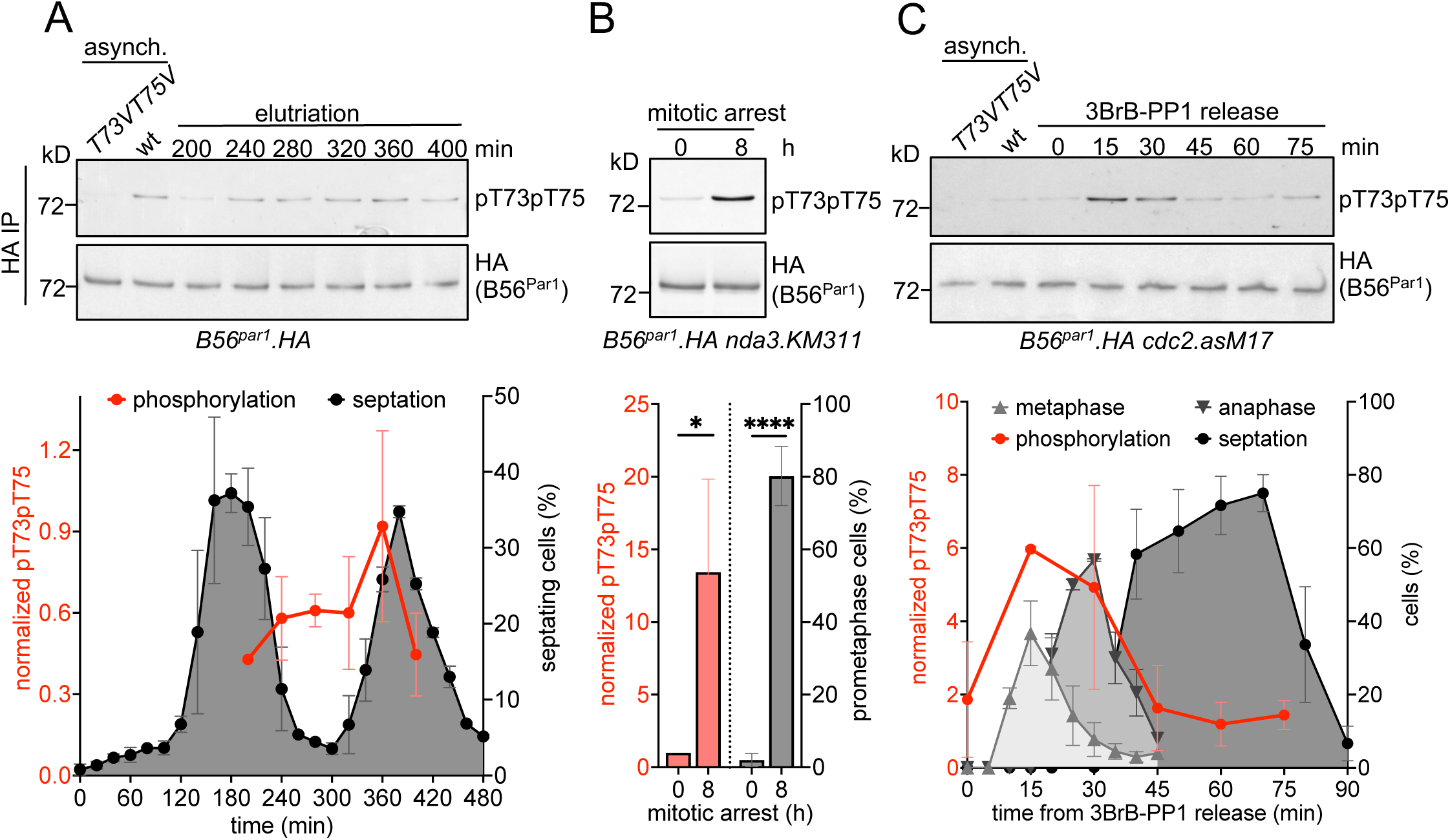
PP2A-B56^Par1^.T73T75 phosphorylation rises in G2 and peaks in mitosis. (**A-C**) B56^Par1^.T73T75 phosphorylation levels in cell cycle-synchronized B56^Par1^.HA cultures. Top, representative immunoblots with samples from asynchronous cultures as controls, bottom, quantification of phosphorylation levels alongside scores of cell cycle stages (means +/- SD, *n*≥2). pT73pT75 levels were normalized to total levels of the B56^Par1^.HA target protein run on a parallel Western blot as sample processing controls, and normalized values were plotted relative to asynchronous wild type, which was set to 1. (**A**) Centrifugal elutriation. Cell cycle progression was monitored by scores of septating cells. (**B**) *nda3.KM311*-mediated mitotic arrest. Arrest was monitored by the accumulation of prometaphase cells that displayed condensed chromosomes. Significance was calculated by ratio paired *t*-test, *P < 0.05 (phosphorylation levels), and unpaired *t*-test, ****P < 0.0001 (mitotically arrested cells). (**C**) Mitotic progression of *cdc2.asM17* cultures upon release from 3BrB-PP1-induced G2 arrest. Cell cycle phases were monitored by scoring for metaphase, anaphase and septating cells. wt, wild type.

To confirm that the B56^Par1^.T73T75 phosphorylation increases in mitosis with a more robust cell synchronization protocol, we took the advantage of the *nda3.KM311* thermosensitive β-tubulin mutant, which arrests cells in mitotic prometaphase in restrictive temperature (Hagan *et al*, 2016c; Hiraoka *et al*, 1984). B56^Par1^.T73T75 phosphorylation was strongly upregulated in the mitotically arrested sample (Fig. 2B).

To further investigate the dynamics of the phosphorylation during mitotic progression, we then followed the phosphorylation kinetics as cells progressed through the different mitotic phases in Adenosine triphosphate (ATP) analogue 3BrB-PP1-synchronized *cdc2.as-M17* cultures (Singh *et al*, 2021). In parallel with sampling for B56^Par1^.T73T75 phosphorylation, we quantified the frequency of metaphase, anaphase and septating cells by staining for β tubulin (mitotic spindles), DAPI (nuclei) and calcofluor (septa). This analysis revealed that T73T75 phosphorylation peaks in metaphase and then declines as cells complete anaphase (Fig. 2C).

Taken together, these results show that B56^Par1^.T73T75 phosphorylation rises in the G2 phase of the cell cycle, peaks in mitosis and declines upon mitotic exit.

### CDK1-Cyclin B phosphorylates PP2A-B56^Par1^.T73T75

The mitotic up-regulation of the phosphorylation prompted us to ask whether it is directly mediated by the major mitotic kinase CDK1-Cyclin B. Sequences surrounding Threonines 73 and 75 match a long [S/T]-P-x-x-[K/R] or short [S/T]-P-[K/R] CDK1 phosphorylation consensus sequence variants, respectively (Fig. 3A). To address whether CDK1-Cyclin B directly phosphorylates these residues, we first asked whether B56^Par1^ serves as a CDK1 phosphorylation substrate. CDK1-Cyclin B^Cdc13^ immuno-precipitated from mitotic *S. pombe* cells using Cks1^Suc1^ affinity directly phosphorylated recombinant B56^Par1^.GST *in vitro*, whilst the phosphorylation was inhibited in the presence of the CDK inhibitor roscovitine (Fig. 3B). This result demonstrates that CDK1-Cyclin B can act as a B56^Par1^ kinase.

**Figure 3.**
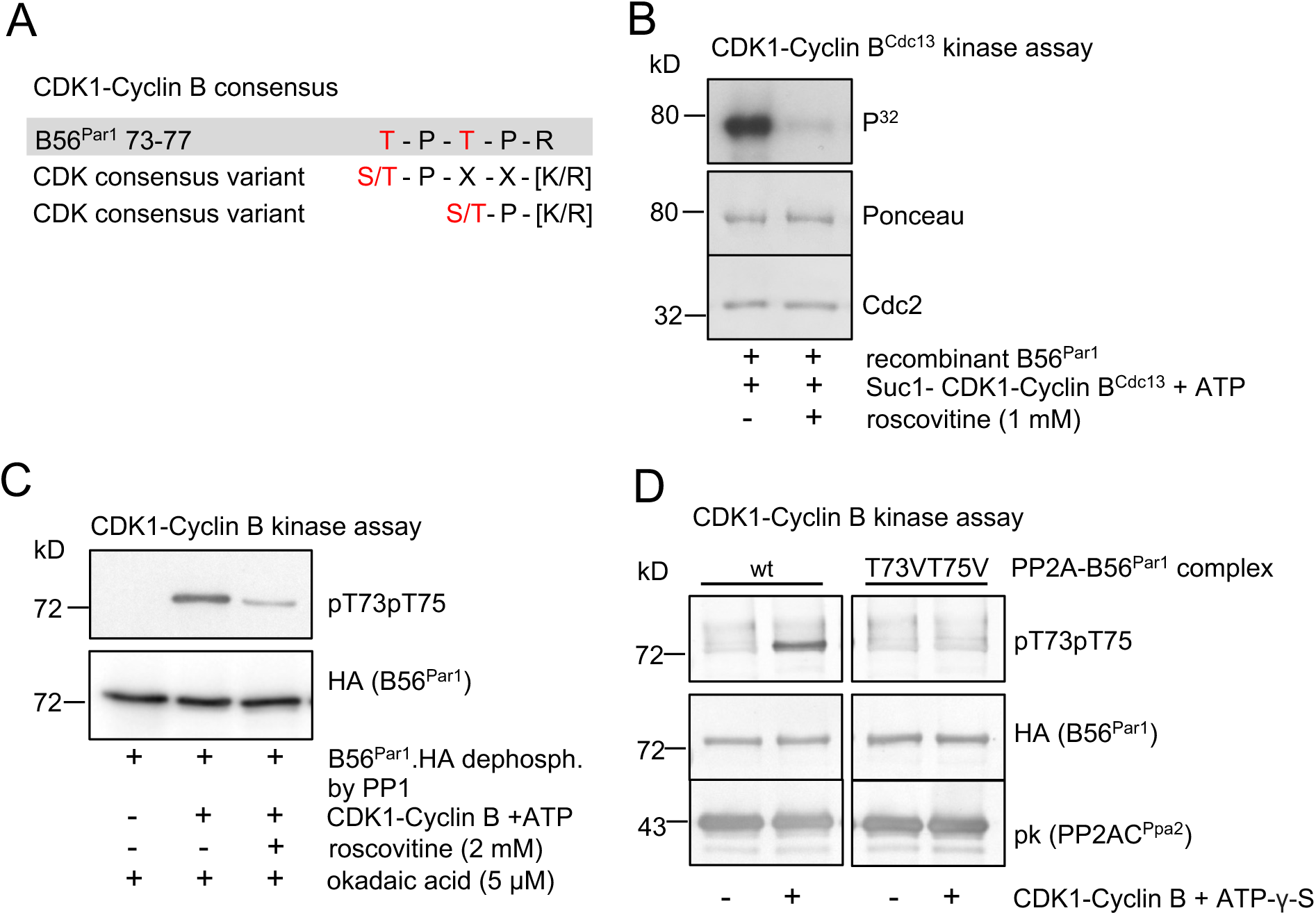
CDK1-Cyclin B phosphorylates PP2A-B56^Par1^.T73T75. (**A**) The sequence surrounding the B56^Par1^.T73T75 phospho-site alongside consensus variant sequences for CDK phosphorylation. (**B-D**) Phosphorylation of B56^Par1^ by CDK1-Cyclin B in *in vitro* kinase assays. (**B**) Recombinant B56^Par1^ was phosphorylated (+/- CDK inhibitor roscovitine) by CDK1-Cyclin B^Cdc13^ precipitated from mitotic *S. pombe* cells by Cks1^Suc1^ affinity. Phosphorylation was detected by autoradiography. Ponceau staining and anti-Cdc2 antibodies against the fission yeast CDK1 were used as loading controls. (**C**) B56^Par1^.HA was HA-immunoprecipitated from asynchronous *S. pombe* cultures under denaturing conditions, de-phosphorylated by PP1 and then phosphorylated by CDK1-Cyclin B (+/- CDK inhibitor) in the presence of the PP1 inhibitor okadaic acid. Phosphorylation was detected by pT73pT75 phospho-specific antibodies and B56^Par1^.HA levels were analysed in a parallel loading on the same Western blot as sample processing controls. (**D**) Endogenous PP2A-B56^Par1^.HA and PP2A-B56^Par1^.T73VT75V.HA complexes were purified from asynchronous cultures and subjected to *thio*-phosphorylation by CDK1-Cyclin B and ATP-γ-S. Phosphorylation was detected by pT73pT75 phospho-specific antibodies and B56^Par1^ and PP2AC^Ppa2^ levels were analysed on a parallel Western blot as sample processing controls. wt, wild type.

We next asked whether CDK1 directly phosphorylated B56^Par1^ isolated from yeast. We immuno-precipitated B56^Par1^.HA from asynchronous *S. pombe* cultures by HA affinity under denaturing conditions, and de-phosphorylated the precipitate with recombinant PP1Cα. The now de-phosphorylated B56^Par1^.HA was then used as a substrate in a CDK1-Cyclin B kinase assay, in the presence of the PP1 and PP2A inhibitor okadaic acid to prevent any dephosphorylations. To determine if CDK1 phosphorylated B56^Par1^.HA on T73T75, we monitored phosphorylation with pT73pT75 phospho-specific antibodies. CDK1-Cyclin B directly phosphorylated B56^Par1^ on T73T75 (Fig. 3C).

The assays shown in Fig. 3B-C utilized an isolated B56^Par1^ molecule as the kinase substrate. In cells however, B56^Par1^ functions as part of the trimeric PP2A-B56^Par1^ complex. To determine whether CDK1-Cyclin B phosphorylates T73T75 of B56^Par1^ when incorporated into the PP2A complex, we purified endogenous PP2A-B56^Par1^ by sequential Tandem Affinity Purification (TAP) tag and HA affinities (Grallert *et al*., 2015). The purified complex was then used as a substrate in the CDK1-Cyclin B kinase assays performed in the presence of a non-hydrolyzable ATP analogue ATP-γ-S to prevent dephosphorylation. Importantly, CDK1-Cyclin B directly phosphorylated T73T75 when B56^Par1^ was purified as part of the endogenous PP2A-B56^Par1^ complex (Fig. 3D). Interestingly, in an assay with hydrolysable ATP, CDK1-Cyclin B could only phosphorylate the complex in the presence of a PP2A inhibitor (nM okadaic acid), indicating that PP2A-B56^Par1^ can auto-dephosphorylate (Appendix Fig. S2 and (Inoue *et al*, 1999; Tanabe *et al*, 1996)).

We conclude that B56^Par1^.T73T75 is a target for CDK1-Cyclin B phosphorylation, and that CDK1-Cyclin B directly phosphorylates PP2A-B56^Par1^ complex on T73T75.

### PP2A-B56^Par1^.T73T75 phosphorylation enhances PP2A-B56^Par1^ activity

Phosphorylations of PP2A subunits have been shown to impact on PP2A complexes function, namely activity or integrity (Ahn *et al*, 2011; Ahn *et al*, 2007; Dodge-Kafka *et al*, 2010; Letourneux *et al*, 2006; Margolis *et al*, 2006; Nasa *et al*., 2020; Shouse *et al*, 2011; Usui *et al*, 1998; Wu *et al*, 2024; Xu & Williams, 2000). We therefore examined the effect of the T73T75 phosphorylation on the PP2A-B56^Par1^ heterotrimer.

Since PP2A-B56 can auto-dephosphorylate (see Appendix Fig. S2 and (Inoue *et al*., 1999; Tanabe *et al*., 1996)), we addressed the impact of T73T75 phosphorylation on PP2A-B56^Par1^ using phospho-mimetic (T73DT75D) and non-phosphorylatable (T73VT75V) variants, which are refractory to dephosphorylation at these sites.

We first asked if mimicking or disabling T73T75 phosphorylation affected the PP2A-B56^Par1^ complexes integrities. PP2A heterotrimer assembly is a stepwise process, where the scaffold (A) subunit and the regulatory (B) subunit form a dimer, which then binds the catalytic (C) subunit to form the final heterotrimer (Janssens & Goris, 2001). To address complex integrity, we therefore monitored the capacity of the regulatory subunit (B56^Par1^) to co-precipitate with the major PP2A catalytic subunit (PP2AC^Ppa2^). Both B56^Par1^ phospho-mimicking (T73DT75D) and non-phosphorylatable (T73VT75V) mutants remained in complex with the catalytic subunit in both asynchronous and analogue-synchronised cultures as cells progressed through the different stages of mitosis (Appendix Fig. S3A-B), confirming that PP2A-B56^Par1^ complex integrity is not compromised by the T73XT75X mutation.

We next asked if mimicking or blocking B56^Par1^.T73T75 phosphorylation impacted on PP2A-B56^Par1^ phosphatase activity. To this end, we purified the major endogenous PP2A-B56^Par1^ complexes [A^Paa1^-B56^Par1^-C^Ppa2^ (Jiang & Hallberg, 2000)] by sequential TAP tag and HA affinities from *S. pombe* cells ((Grallert *et al*., 2015); Appendix Fig. S4A and Supplementary Data 2). To measure the purified complexes activities, we adopted a phosphatase assay developed for human PP2A-B56 (Hein *et al*, 2021; Kruse *et al*., 2020). The substrate in this assay is an engineered Cdc20 fragment fused to a PP2A-B56 docking site motif LDTIQEEE (LxxIxE) and phosphorylated on N-terminal Threonine-Proline sites by Cdk1-Cyclin B (Fig 4A). The assay was optimized to measure substrate dephosphorylation over a period of up to two hours (Appendix Fig. S4B-C).

**Figure 4.**
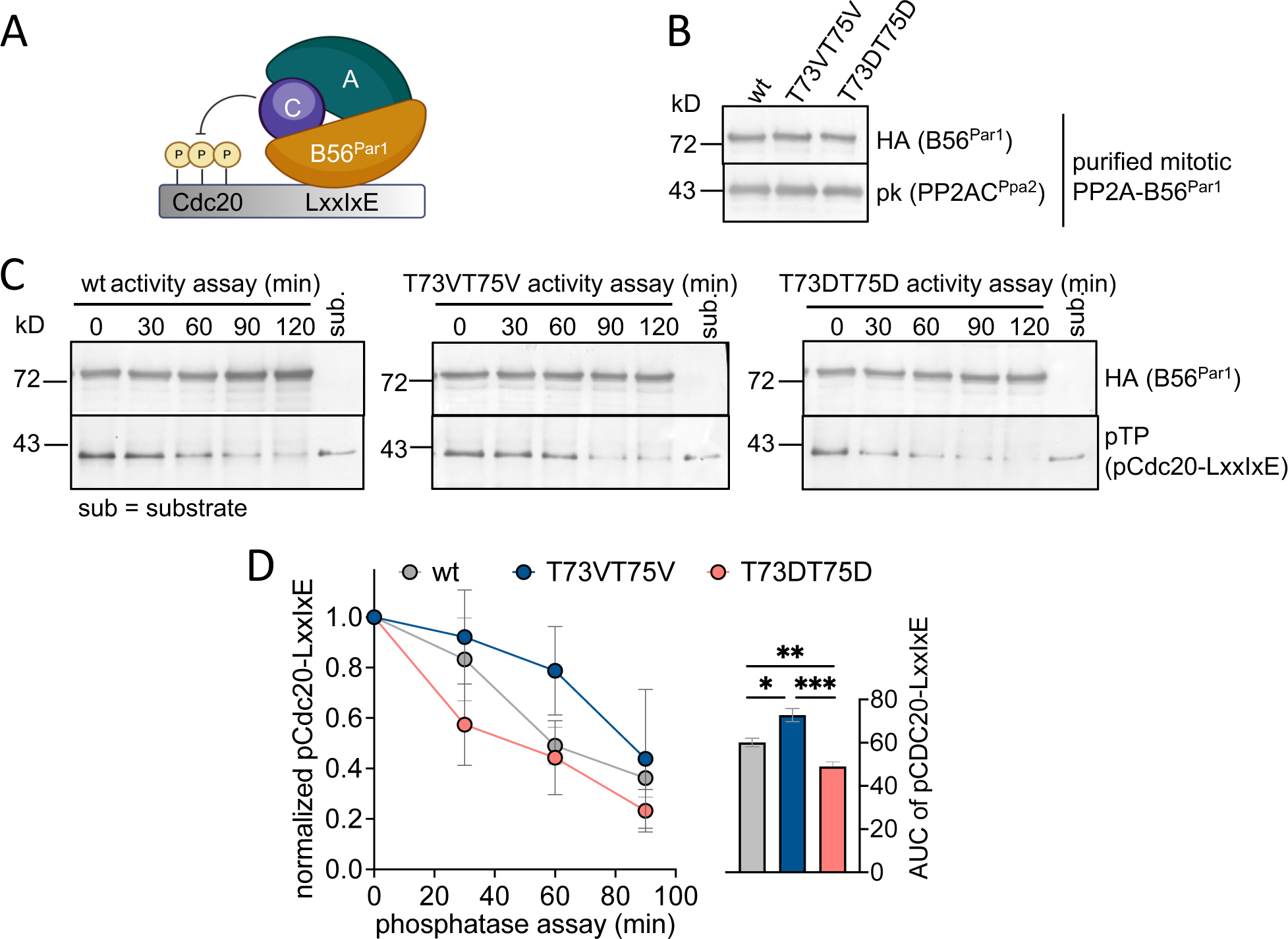
Mimicking PP2A-B56^Par1^.T73T75 phosphorylation boosts while disabling blocks PP2A-B56^Par1^ complex activity. (**A-D**) Activity assay of purified endogenous PP2A-B56^Par1^ complexes. (**A**) Schematic of the activity assay. The purified PP2A-B56^Par1^ complex activity was measured *in vitro* by monitoring the rate of dephosphorylation of a Cdc20 substrate harboring the PP2A-B56 LxxIxE docking motif (Kruse *et al*., 2020). (**B-C**) Equal amounts of purified mitotic PP2A-B56^Par1^ complexes variants (**B**) were mixed with phosphorylated Cdc20-LxxIxE substrate and dephosphorylation was monitored for 120 min with aliquots withdrawn every 30 min (**C**). (**D**) Quantification of activity measured by level of substrate dephosphorylation (means +/-SD, *n*=4). pCdc20-LxxIxE levels were normalized against loading control (B56^Par1^.HA), and normalized values were plotted relative to time point 0 min, which was set to 1. Area under the curve (AUC) significance was calculated by unpaired *t*-test, *P < 0.05, **P < 0.01, ***P < 0.001. T73VT75V, B56^Par1^.T73VT75V.HA complex; T73DT75D, B56^Par1^.T73DT75D.HA complex; wt, wild-type complex.

T73T75 phosphorylation is maximal in mitosis (see Fig. 2A-C), which prompted us to determine the impact of the T73XT75X mutations on the activity of PP2A-B56^Par1^ complexes purified from mitotic cells. We arrested cells expressing endogenous T73XT75X complexes variants in mitosis by *nda3.KM311-*mediated arrest (Appendix Fig. S5A) and purified the PP2A-B56^Par1^ complexes by sequential TAP tag – HA tag affinities (Fig 4B). We then measured the purified mitotic complexes activity towards phosphorylated Cdc20-LxxIxE. The phospho-mimetic variant of the complex (T73DT75D) was able to dephosphorylate the substrate with a faster kinetics compared to wild type, while the non-phosphorylatable variant (T73VT75V) was less effective (Fig. 4C-D). The activity was blocked by a PP2A inhibitor (Appendix Fig S5B-C). A similar trend in activities of the T73DT75D and T73VT75V variants was observed with purifications from asynchronous cultures (Appendix Fig. S6A-C). These results indicate that phosphorylation of B56^Par1^.T73T75 increases the PP2A-B56^Par1^ complex activity, whilst disabling the phosphorylation blocks activity.

### PP2A-B56^Par1^.T73T75 phosphorylation is not required for mitotic progression

As B56^Par1^.T73T75 phosphorylation increases at G2/M transition and declines upon mitotic exit (see Fig. 2A-C), we next examined the requirement of the B56^Par1^.T73T75 phosphorylation for mitotic progression. To this end, we generated endogenous un-tagged *B56^par1^*non-phosphorylatable (*T73VT75V*) and phospho-mimetic (*T73DT75D*) mutants and assessed their phenotypes.

PP2A-B56 has been shown to regulate events at metaphase/anaphase transition in both fission yeast (Deng *et al*., 2022; Grallert *et al*., 2015; Kitajima *et al*., 2006; Tanabe *et al*, 2001) and higher Eukaryotes (Espert *et al*, 2014; Foley *et al*, 2011; Fujimitsu & Yamano, 2020; Kruse *et al*., 2013; Nijenhuis *et al*., 2014; Quiniou *et al*, 2025; Ueki *et al*, 2021). In fission yeast, B56^Par1^ is also required for cytokinesis (Chrupcala & Moseley, 2025). To ask if the phosphorylation impacts on mitotic progression, we monitored the variants’ progression through mitosis and cytokinesis in analogue-synchronized cultures. Both *T73DT75D* and *T73VT75V* mutants progressed normally through the different mitotic phases and septation (Fig. 5A), indicating that the phosphorylation was not required for mitotic progression and cytokinesis.

**Figure 5.**
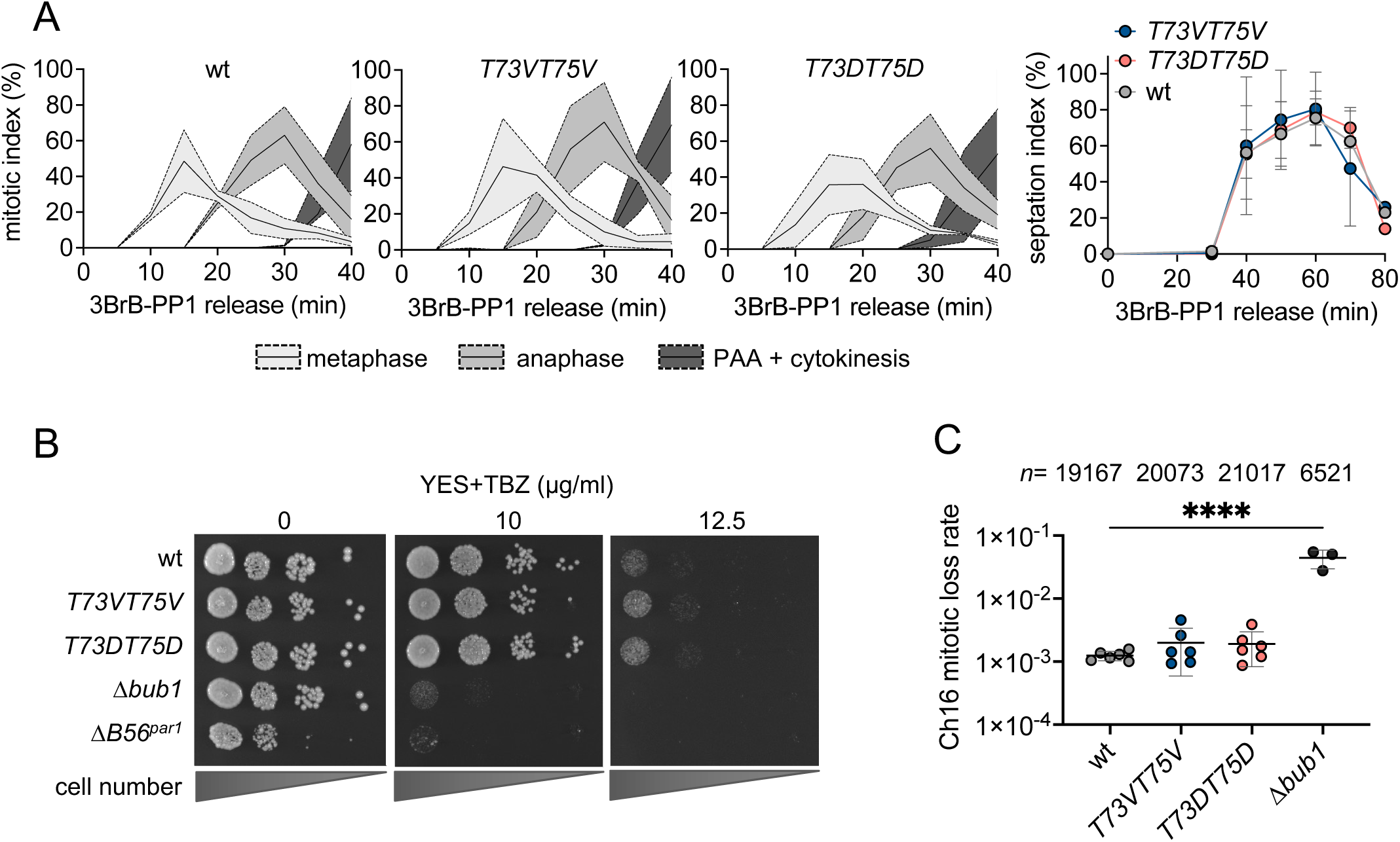
PP2A-B56^Par1^.T73T75 phosphorylation is not required for progression through mitosis. (**A**) Mitotic index (left) and septation index (right) of *cdc2.as-M17 B56^par1^.T73XT75X* cultures upon release from 3BrB-PP1-induced G2 arrest (means +/- SD, *n*=2). (**B**) Spot test of serial dilutions of asynchronous yeast cultures assessing TBZ sensitivity. The *Δbub1* strain lacking the essential SAC component Bub1 (Bernard *et al*, 1998) was used as a SAC-deficient control. (**C**) Minichromosome (Ch16) mitotic loss assay. The *Δbub1* strain was used as a chromosome segregation fidelity - deficient control. Significance was calculated by a one-way ANOVA followed by Dunnett’s multiple comparisons test, ****P < 0.0001.

Although not a key player, PP2A-B56^Par1^ has been shown to function in spindle assembly checkpoint (SAC) silencing (Deng *et al*., 2022). Mutations in genes required for chromosome segregation, such as kinetochore- or SAC-related genes, sensitize *S. pombe* cells to a microtubule-destabilizing drug, Thiabendazole (TBZ; (Akera *et al*, 2015; Bai *et al*, 2022)). Δ*B56^par1^* cells show TBZ sensitivity (Tanabe *et al*., 2001). To test the role of the T73T75 phosphorylation in SAC control, we assessed the phospho-mutants’ growth in the presence of TBZ. Increasing concentrations of TBZ had no impact on growth of *B56^par1^.T73XT75X* (Fig. 5B), suggesting a normal SAC function. In addition to TBZ sensitivity, *ΔB56^par1^*cells experience chromosome segregation errors (Grallert *et al*., 2015). We therefore addressed the chromosome segregation fidelity of *T73XT75X* mutants in an assay that measures the stability of an artificial mini-chromosome Ch16 (Niwa, 2018). Ch16 stability was not affected by neither of the *B56^par1^.T73XT75X* phospho-mutations (Fig. 5C). These results demonstrate that under our experimental conditions, T73T75 phosphorylation has no impact on SAC control or chromosome segregation fidelity.

Taken together, these results indicate that T73T75 phosphorylation is obsolete once cells are in mitosis.

### Blocking PP2A-B56^Par1^.T73T75 phosphorylation leads to premature division at a smaller cell size

We next investigated the role of T73T75 phosphorylation in G2/M transition. *ΔB56^par1^* cells divide on average with a shorter cell size than wild type, with a wider distribution of sizes (Fig. 6A-B and (Grallert *et al*., 2015; Tanabe *et al*., 2001)), implying that the activity of PP2A-B56^Par1^ is required for cell size control at division. Additionally, *ΔB56^par1^*cells exhibit morphological and septation defects (arrowheads in Fig. 6B), consistent with PP2A-B56^Par1^ role in cell polarity and cytokinesis (Chica *et al*, 2022; Chrupcala & Moseley, 2025; Jiang & Hallberg, 2000, 2001; Tanabe *et al*., 2001). Intriguingly, cells expressing the non-phosphorylatable *B56^par1^.T73VT75V* mutant divided at an average cell length approximately 2µm shorter than wild type, corresponding to a 15% reduction in cell length (Fig. 6A-B). To corroborate this finding, we generated an independent non-phosphorylatable variant, *B56^par1^.T73AT75A*, and observed a similar small-size phenotype. These results show that blocking T73T75 phosphorylation leads to division at a reduced cell size. In line with that, phospho-mimetic *B56^par1^.T73DT75D* maintained wild-type length at division (Fig. 6A-B). Neither *T73XT75X* mutant displayed morphological or septation defects seen in *ΔB56^par1^*. In media with glucose and a rich nitrogen source, such as EMM2 used in our study, *S. pombe* cells only grow in G2 phase of the cell cycle (due to a cryptic G1 cell size checkpoint) and arrest growth as they commit to mitosis with a defined cell size. Cell size at division is therefore a function of mitotic entry timing (Mitchison & Nurse, 1985; Moreno & Nurse, 1994; Petersen & Nurse, 2007).

**Figure 6.**
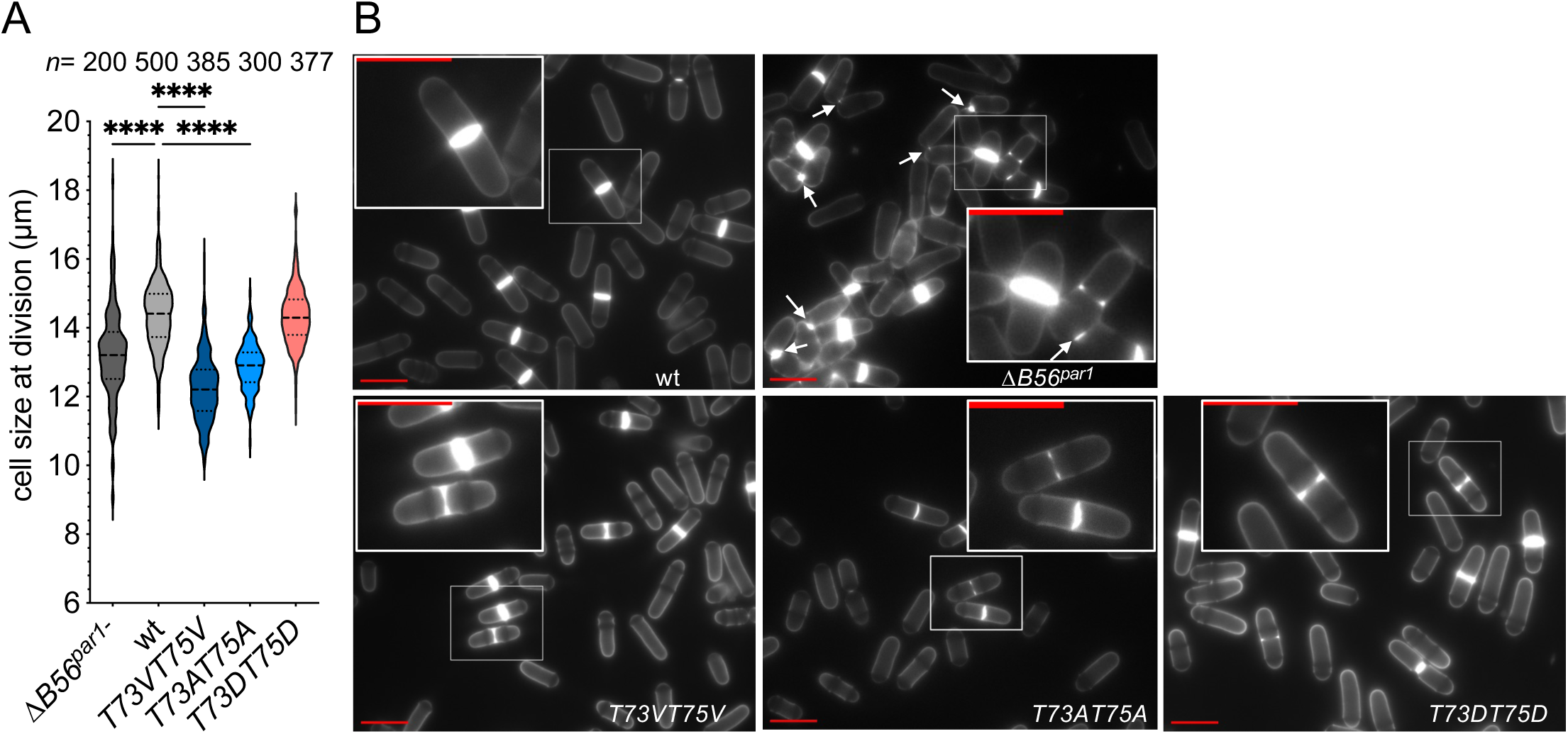
Blocking PP2A-B56^Par1^.T73T75 phosphorylation induces premature division at a reduced cell size. (**A-B**) Cell size at division of asynchronous unperturbed cultures of endogenous B56^Par1^.T73XT75X mutants. (**A**) Quantification of cell division sizes. *ΔB56^par1^*and phospho-blocking *B56^par1^.T73VT75V* and *B56^par1^.T73AT75A* divide at a reduced cell size. Significance was calculated by Kruskal-Wallis’s test followed by Dunn’s multiple comparisons test. **P < 0.01, ****P < 0.0001. (**B**) Representative images with inset magnifications of calcofluor-stained cultures. Arrowheads mark septation defects present in *ΔB56^par1^*. Scale bar, 10 µm.

We conclude that B56^Par1^.T73T75 phosphorylation is required to maintain cell size control at mitotic entry in unperturbed cycles. Disabling T73T75 phosphorylation leads to premature entry into mitosis with a reduced cell size.

## Discussion

We initiated the present study by searching for PP1^Dis2^ target sites on B56^Par1^, which led to the identification of Proline-directed regulatory sites T73T75 in the N-terminal arm of B56^Par1^. Previously, we demonstrated that PP1^Dis2^ directly regulates PP2A-B56^Par1^ during mitotic exit through docking to G/SILK/R and K/RxVxF motifs on B56^Par1^ (Grallert *et al*., 2015). In contrast, the regulatory mechanism described here is independent of this docking interaction (Appendix Fig. S1C-D), raising the possibility that PP1^Dis2^ interacts with B56^Par1^ through an alternative mechanism or exerts its effect indirectly. The latter is supported by observations that both PP1 and PP2A-B56 dock to regulatory or intermediate targets that subsequently recruit them to their substrates (Bollen *et al*., 2010; Suijkerbuijk *et al*, 2012).

Analysis of endogenous T73T75 phospho-mutants revealed an outstanding cell size phenotype. Blocking B56^Par1^.T73T75 phosphorylation reduced cell length at division in unperturbed cycles from 14 to 12 µm (Fig. 6A), dramatically altering the surface are-to-volume ratio. This 2µm cell size reduction demonstrated that blocking the phosphorylation prematurely unlocked the amplification of the CDK1/Cdc25/Wee1 feedback loop, which drives mitotic onset (Ferrell, 2008; Hagan & Grallert, 2013; Mitchison & Nurse, 1985; Moreno & Nurse, 1994; Petersen & Nurse, 2007).

The B56^Par1^.T73T75 phosphorylation is directly mediated by CDK1-Cyclin B (Fig. 3A-D) and phospho-mimicking mutation increases PP2A-B56^Par1^ phosphatase activity while non-phosphorylatable blocks activity (Fig. 4A-C and Appendix Fig. S6A-C). We hypothesize that by phosphorylating T73T75, the rising CDK1-Cyclin B activity concurrently activates the counteracting PP2A-B56^Par1^ phosphatase, limiting the net accumulation of substrates phosphorylations. Blocking T73T75 phosphorylation reduces PP2A-B56 activity, leading to increased substrates phosphorylations, accelerated mitotic onset, and division at an approximately 2µm smaller cell size. A similar concept where phosphatase activity rises concurrently with a counteracting kinase during G2 has been proposed for PP2A-B55 in *Xenopus* (Kamenz et al., 2021).

Mitosis is triggered when the CDK/Cdc25/Wee1 feedback loop signalling at the spindle pole body (SPB) overcomes inhibitory signals from counteracting phosphatases and from Wee1 signalling emanating from the cell equator (Allard *et al*, 2018; Hagan & Grallert, 2013; Kapadia & Nurse, 2025). Curiously, B56^Par1^ is found both on the SPB and in equatorial cortical nodes (Chrupcala & Moseley, 2025; Le Goff *et al*, 2001). The premature mitosis of the lower-activity phospho-blocking B56^Par1^.T73T75 mutants could result from activation of Cdc25, inhibition of Wee1, or a combination of the two. Interestingly, in *Xenopus* and human cells, PP2A-B56 has been shown to directly dephosphorylate and inactivate Cdc25, as part of the DNA damage checkpoint signalling in interphase (Margolis *et al*., 2006) and at mitotic exit (Forester *et al*, 2007). In fission yeast, CDK-mediated phosphorylations of Cdc25, including phosphorylations of T104, S204 and T379, were identified as PP2A-B56 targets upon acute PP2A-B56 depletion in mitosis (Zeisner *et al*., 2025). Additionally, the Nurse lab analysis identified regulatory phosphorylation sites of the Cdc25 phosphatase CDC14^Flp1^, including inhibitory T453 phosphorylation, as PP2A-B56 substrates (Wolfe & Gould, 2004; Wolfe *et al*, 2006; Zeisner *et al*., 2025). It remains to be determined whether PP2A-B56 prevents premature mitotic entry by inhibiting Cdc25, either directly by dephosphorylation of CDK sites, or indirectly through dephosphorylation and activation of CDC14^Flp1^, or through a combination of both mechanisms.

The fission yeast SPB, like the human centrosome, integrates signals from mitotic regulators (Arquint *et al*, 2014; Chan *et al*, 2017; Hagan & Grallert, 2013; Lin *et al*, 2022). Other CDK-opposed phosphatases have been shown to regulate SPB activation and modulate Cdc25 or Wee1 signalling. PP1^Dis2^ recruitment to SPB governs the timing of the SPB activation in G2 (Basu *et al*., 2022; Grallert *et al*., 2013a). CDC14^Flp1^ eviction from the SPB at mitotic commitment reduces the local threshold for CDK1-Cyclin B activation on the SPB (Chan *et al*., 2017; Wolfe & Gould, 2004). PP2A-B55^Pab1^ maintains low CDK1 activity by dephosphorylating Cdc25 and Wee1 that inhibits and activates the respective CDK1 regulators (Chica *et al*., 2016; Lucena *et al*, 2017), while Wee1 localized to cortical ‘nodes’ that form a band around the cell equator blocks propagation of feedback loop signalling from the SPB (Allard *et al*., 2018). The above discussion, together with our observation that PP2A-B56^Par1^ mutant cells divide at a 2µm smaller cell size, supports a tentative model in which the SPB integrates diverse CDK-opposed phosphatase signalling prior to mitosis. We propose that phosphatase inputs converging at the SPB are integrated into a coherent signal that is amplified throughout the cell to regulate mitotic entry. Although speculative, this model provides a useful framework for future experimental investigations.

To our knowledge, this study is the first to demonstrate that PP2A-B56 regulation controls mitotic entry timing during unperturbed cycles. The Threonine-Proline module is conserved in analogous regions of B56 orthologs, including humans (Fig. 1D). Given the broad conservation of cell cycle control by CDK-opposed phosphatases from *S. pombe* to higher Eukaryotes, it is likely that similar regulatory mechanisms operate in human cells.

## Materials and methods

### Yeast cell culture

*S. pombe* strains used for this study are listed in Appendix Table S1. For standard growth and cell cycle synchronization assays, cells from -70°C glycerol stocks were streaked on YES+ amino acids (AA) medium (YES supplemented with 250 mg/L Adenine, Histidine, Leucine, Uracil and Lysine, see Reagents and Tools Table) and then grown in liquid Edinburgh minimal media 2 (EMM2, Appendix Table S2) without supplements (unless otherwise specified) at 25°C in the log phase for 48h until the density of 2x10^6^ cells/ml. For cell size measurements experiments, cells were grown in EMM2 without supplements at 32°C in the log phase for 48h until the density of 2x10^6^ cells/ml.

Centrifugal elutriation was performed as described (Hagan *et al*., 2016a). For ATP-analogue synchronizations of mitotic entry, *cdc2.as-M17* cultures were arrested in G2 for 4h 15min by 2µM ATP-analogue 3Br-BPP1 and then filtered into a fresh media to release from inhibition (Singh *et al*., 2021). *nda3.KM311*–mediated arrest was performed in YES for 8h at 20°C (Hagan *et al*., 2016c).

For large-scale affinity purifications, cells were inoculated in YES+AA medium overnight and then diluted in 2x concentrated YES+AA and grown for 48h up to the density of 1x10^8^ cells/ml.

### Large scale affinity purification for Quantitative Mass Spectrometry

2x10^11^ cells/sample grown were processed according to a standard protocol (Grallert & Hagan, 2017a). Briefly, cells were cryo-milled under liquid nitrogen in a SPEX Sample Prep 6875D freezer mill (8 cycles 2-2-2 with impact rate of 14). Solubilization was achieved by the addition of 6% SDS and immunoprecipitation was performed under denaturing conditions (Grallert & Hagan, 2017b) with HA-affinity (Pierce anti-HA beads) in immunoprecipitation (IP) buffer (50 mM Hepes pH 7.5, 1 mM Ethylenediaminetetraacetic acid [EDTA], 100 mM NaCl, 40 mM Na-β-glycerophosphate, 2 mM Na_3_VO_4_, 50 mM Sodium fluoride [NaF], 0.1% NP40, 1 mM Phenylmethylsulfonyl fluoride [PMSF], cOmplete™ EDTA-free Protease Inhibitor Cocktail) for 1h at 4°C. HA-precipitates were resolved in a 4-12% 2-[Bis(2-hydroxyethyl)amino]-2-(hydroxymethyl)propane-1,3-diol (Bis-Tris) gel and stained with Coomassie Brilliant Blue G-Colloidal concentrate.

### Mass Spectrometry

Protein bands were excised from gel destained with three 10min washes in 1 ml high-performance liquid chromatography (HPLC) grade water with 40% acetonitrile and 200mM ammonium bicarbonate. Samples were then cleaned with a 10min wash with 1ml HPLC grade water followed by dehydration with 1 ml of acetonitrile. This cycle of hydration-dehydration was performed a total of 3 times followed by vacuum centrifugation. Protein bands were digested with 20 ng sequencing-grade trypsin in 100 μl 40 mM ammonium bicarbonate with 9% (v/v) acetonitrile at 37°C for 18h. The peptides were separated using an RSLCnano Ultra-High Performance Liquid Chromatography (UPLC) system using a Thermo pepmap C18 EasySpray column (75 μm inner diameter, 2 μm, 50 cm) with a gradient of 1–25% (v/v) of acetonitrile with 0.1% formic acid over 30min at a flow rate of 200 nl/min. The Orbitrap Fusion Mass Spectrometer was operated in data-dependent mode, where the survey scan was performed at a nominal resolution of 120,000 (at m/z 200) in the Orbitrap analyzer over an m/z range of 350–1200. The instrument performed as many Higher-Energy Collisional Dissociation (HCD) second-stage mass spectrometry (MS2) scans as possible in a total cycle time of 1.1s. HCD was performed at 28% normalized collision energy and the MS2 spectra were collected in the linear ion trap. For protein identification and phospho-mapping, Mascot Generic Format datasets (MGFs) were generated in Mascot Distiller followed by a database search using Mascot (Matrix Science). Mascot search results were imported into Scaffold and Scaffold PTM. Label-free quantification of phoshopeptides was achieved with Progenesis LC-MS.

### Genetic manipulations

Cloning primers are listed in Appendix Table S3. To generate endogenous *B56^par1^.T73T75* phospho-mutants, mutations encoding T73XT75X variants were introduced via site-directed mutagenesis into a Topoisomerase based cloning (TOPO) vector harboring B56^Par1^ open reading frame (ORF) flanked by 100 bp 5’ and 3’ untranslated regions (UTRs) (Grallert *et al*., 2015). The clones were transformed into *S. pombe ΔB56^par1^* host according to standard procedures (Fennessy *et al*, 2014), selected by 5-Fluoroorotic acid (5-FOA) resistance and mutation confirmed by DNA sequencing. HA-tagging of endogenous *B56^par1^* was performed as previously described (Bahler *et al*, 1998).

To generate *pk.PP2AC^ppa2^*, the *PP2AC^ppa2^*ORF flanked by 100 bp 3’ and 5’ UTRs was amplified by PCR and inserted into a TOPO vector. N-terminal pk tag and Hygromycin-resistant allele *hphMX6* were inserted into the resulting plasmid by Gibson assembly (Gibson *et al*, 2009). The *pk.ppa2:hpHMX6* cassette was then transformed into *S. pombe ΔPP2AC^ppa2^*host according to standard procedures (Fennessy *et al*., 2014), clones selected by 5-FOA resistance and insertion confirmed by DNA sequencing.

### Small scale HA-affinity purification

2x10^8^ cells (6x10^8^ for *B56^par1^.PDSN*) were harvested by centrifugation (3500g for 2 min), washed in STOP buffer (50 mM NaF, 10 mM EDTA, 0.9% NaCl, 1 mM Sodium azide [NaN_3_]) at 4°C and pellets were stored at -70°C until processing. HA-affinity purification was performed as previously described under denaturing or native conditions (Grallert & Hagan, 2017b). Briefly, cell pellets were disrupted using a multi-beads shocker and immunoprecipitation was performed with Pierce HA beads in IP buffer for 1h at 4°C.

### PP1^Dis2^ phosphatase assay

B56^Par1^.HA substrate was precipitated from wild-type cells by small scale denaturing IP (Grallert & Hagan, 2017b). The HA-beads-bound precipitates were then washed in Protein MetalloPhosphatases (PMP) buffer (50 mM HEPES pH 7.5, 100 mM NaCl, 2 mM Dithiothreitol [DTT], 0.01% Brij-35, 1mM MnCl_2_) and incubated in 10 µl PMP buffer with 1.38 µg PP1C⍺ without or with inhibitor (5 µM okadaic acid) for 1h at 30°C.

### Microscopy

Cell cycle progression was tracked by scoring the percentage of septating cells with calcofluor and/or monitoring chromatin condensation with 4′,6-diamidino-2-phenylindole (DAPI) according to standard procedures (Hagan, 2016a). Mitotic phases were monitored by combined DAPI staining and anti-tubulin immunofluorescence with Tat1 tissue culture supernatant monoclonal antibody (1:80, a gift from K. Gull (Woods *et al*, 1989)), using established procedures (Chan *et al*., 2017; Hagan, 2016b). Images were obtained using a ZEISS Axio Imager.M2 microscope with an α Plan-FLUAR 100x/1.45 oil objective (ZEISS) and ZEISS Axiocam 705 mono camera and processed with Fiji.

### Recombinant protein expression and purification

*B56^par1^* coding sequence (CDS; custom made by GeneArt, Life Technologies) was cloned in pETDuet-1 by Sac*I* and Not*I* restriction digest and ligation and His.B56^Par1^ was expressed in *E. coli* BL21-CodonPlus. pRK753 plasmid harboring *MBP.His.TEV* (Tobacco Etch Virus protease; a gift from M. Lowe) was expressed in *E. coli* BL21(DE3)-RIL. *Cdc20^49-78^* fragment fused with “superbinder” LxxIxE peptide LDTIQEE (Kruse *et al*., 2020) was cloned by Nco*I* BamH*I* restriction digest and ligation into pET41 and Cdc20-LxxIxE.His was expressed in *E. coli* BL21-CodonPlus. Protein expression and purification were performed according to standard protocols with HisPur^Tm^ nitrilotriacetic acid (Ni-NTA) agarose under native conditions.

### PP2A-B56^Par1^ complex purification

PP2A-B56^Par1^ complexes harboring the major catalytic subunit PP2AC^Ppa2^ were purified by sequential TAP tag and HA-affinities (Grallert *et al*., 2015). Briefly, 1x10^11^ cell pellets were cryo-milled under liquid nitrogen in a SPEX Sample Prep 6875D freezer mill (8 cycles 2-2-2 with impact rate of 14). Immunoglobulin G (IgG) affinity was performed with 0.5 ml IgG Sepharose 6 fast flow beads in IP buffer without phosphatase inhibitors (50 mM Hepes pH 7.5, 1 mM EDTA, 100 mM NaCl, 1 mM PMSF, 0.1% NP40, cOmplete™ EDTA-free Protease Inhibitor Cocktail) at 4°C for 1h. Bound PP2A complexes were washed twice with IP buffer without phosphatase inhibitors and twice with TEV buffer (10 mM Tris pH 8.0, 150 mM NaCl, 0.1% NP40, 0.5 mM EDTA, 1 mM DTT, cOmplete™ EDTA-free Protease Inhibitor Cocktail) by gravity flow in Econo-Pac Chromatography columns. Columns were then capped at both ends and incubated with 50 µg TEV protease and 500 µl TEV buffer at 25°C for 2h while gently shaking at 100 rpm. Flow-through was then collected and topped with IP buffer without phosphatase inhibitors up to 7 ml and incubated O/N at 4°C with 125 µl Pierce A/G beads covalently coated with monoclonal 12CA5 HA antibody. The HA beads-bound PP2A-B56^Par1^ complexes were then washed 4x with IP buffer without phosphatase inhibitors at 4°C and either immediately used for enzymatic assays, or eluted at 30°C for 1h at 100 rpm with 1 mg/ml HA peptide in elution buffer (20 mM Tris pH 7.4, 150 mM NaCl, 1 mM MnCl_2_, cOmplete™ EDTA-free Protease Inhibitor Cocktail) for protein identification by Mass Spectrometry.

### PP2A-B56^Par1^ activity assay

Phosphatase assays were performed with purified PP2A-B56^Par1^ complexes as previously described (Kruse *et al*., 2020). Cdc20-LxxIxE fragment (150 µg) was phosphorylated with 0.7 ug recombinant Cdk1-Cyclin B kinase and 100 µM ATP in 200 µl kinase assay buffer (KAB; 50 mM Tris pH 7.4, 10 mM MgCl_2_, 1 mM DTT, cOmplete™ EDTA-free Protease Inhibitor Cocktail) for 30 min at 30°C. The reaction was stopped with the addition of 1 mM roscovitine, buffer was then exchanged for phosphatase buffer (50 mM Tris pH 7.4, 150 mM NaCl, 1 mM MnCl_2_, 1 mM DTT, 0.1% Igepal, cOmplete™ EDTA-free Protease Inhibitor Cocktail, 1 mM roscovitine) + 25% glycerol using ZebaSpin desalting columns and the phosphorylated Cdc20-LxxIxE fragment was stored in 20 µl aliquots at -80°C at the concentration of 0.4 µg/µl.

Purified PP2A-B56^Par1^ complexes bound to HA-beads were washed with phosphatase buffer in non-stick tubes on ice and then mixed with 1.2 µg of the phosphorylated Cdc20-LxxIxE substrate in 50 µl phosphatase buffer. Reaction was incubated on a rotator wheel for up to 120 min at 30°C with samples withdrawn at intervals. Reaction was stopped by the addition of Lithium dodecyl sulfate (LDS) sample buffer (NuPage) + 25% ß-mercaptoethanol for 10 min at 70°C.

### CDK1-Cyclin B kinase assay

For *S. pombe* CDK1-Cyclin B^Cdc13^ kinase assay, *cdc25.22* cultures grown at permissive temperature (25°C) in EMM2 + AA (EMM2 supplemented with 250 mg/L Adenine, Histidine, Leucine, Uracil and Lysine) were arrested at G2/M transition by incubation at 37°C for 4h 30 min and then released into permissive 25°C (Hagan *et al*, 2016b). 20 min later, 2x10^8^ cells were harvested by centrifugation (3500g, 2 min) and CDK1-Cyclin B^Cdc13^ was precipitated by small scale CKS1^Suc1^ affinity as previously described (Booher *et al*, 1989; Singh *et al*., 2021). Recombinant His.B56^Par1^ substrate (1.2 µg) was added to 20 µl CKS1^Suc1^ beads with immobilized CDK1-Cyclin B^Cdc13^ and kinase reaction was performed according to protocol (Booher *et al*., 1989) with ATP-γ-P^32^ for 10 min at 30°C without or with inhibitor (1 mM roscovitine).

Recombinant human Cdk1-Cyclin B kinase assays were performed with B56^Par1^ substrates (B56^Par1^.HA subunit or PP2A-B56^Par1^.HA complex) isolated from *S. pombe*. B56^Par1^.HA was HA-affinity purified under denaturing conditions from *S. pombe* asynchronous cultures, *in vitro* dephosphorylated with PP1C⍺, washed twice with KAB and resuspended in 10 µl KAB + 5 µM okadaic acid. The PP2A-B56^Par1^.HA complexes harboring the major pk.PP2AC^Ppa2^ catalytic subunit and bound to HA-beads were washed twice with KAB and resuspended in 10 µl KAB ± 20 nM okadaic acid. 0.5 µg of active recombinant Cdk1-Cyclin B kinase was mixed with the HA beads-bound B56^Par1^ or PP2A-B56^Par1^, respectively, in the presence of 100 µM ATP or ATP-γ-S without or with inhibitor (roscovitine, 2 mM) and okadaic acid (5 µM) as indicated, and the reaction was incubated for 30 min at 30°C.

### Western blotting

Proteins in LDS sample buffer + 25% ß-mercaptoethanol were heated for 10 min at 70°C, separated on a 4-12% Bis-Tris gel and transferred onto a nitrocellulose membrane. Ponceau staining was used as indicated to monitor total protein levels prior to probing with antibodies. Polyclonal antibodies raised against the dual-phosphorylated pT73pT75 were custom-generated and purified by Eurogentec. The following dilutions of antibodies were used in this study: anti-pk clone SV5-Pk1 (1:500), tissue culture supernatant anti-HA clone 12CA5 (1:100), anti-Cdc2 (1:2000), anti-phospho-Threonine-Proline (1:2000), anti-phospho-B56^Par1^.T73T75 (1:1000). Horseradish peroxidase (HRP) or alkaline phosphatase-coupled secondary antibodies and chromogenic detection, chemiluminescence or autoradiography were used to detect the signals. In case of re-probing, the membrane was first incubated in a stripping buffer (0.2M glycine, 0.05% Tween, 0.7% ß-mercaptoethanol, pH 2.5) for 30 min at 60°C before probing with a second set of antibodies.

### TBZ sensitivity assay

Cultures were grown exponentially for 48h in YES+AA medium up to 1x10^6^ cells/ml and serial 10x dilutions were spotted onto YES+AA plates containing increasing concentrations of TBZ. Colony formation was monitored for 4-6 days at 25°C.

### Ch16 mitotic loss rate

Ch16 mitotic loss assays were performed as previously described (Niwa, 2018). Strains harboring artificial Ch16 were grown exponentially for 48h in Edinburgh Minimal Medium Glutamate (EMMG; (Petersen & Russell, 2016)) without supplements up to the density of 1x10^6^ cells/ml and aliquots of 10-100 µl were plated on EMMG plates supplemented with 5 mg/ml Adenine. Frequency of Ch16 loss was scored by monitoring the frequency of half sector red colonies after 6 days at 25°C.

## Authorship and Acknowledgments

Author contributions: L.H.: Conceptualization, Data curation, Formal analysis, Investigation, Methodology, Project administration, Supervision, Validation, Visualization, Writing-original draft, Writing – review & editing. Y.C.: Investigation (Mass spectrometry). D.L.S.: Formal analysis and Data curation (Mass spectrometry). J.P.: Supervision, Writing - review & editing. I.M.H.: Conceptualization, Funding acquisition, Methodology, Project administration, Supervision, Writing – review & editing.

We thank Jon Pines for advice during the last year of the study, Ray O’Keefe, Steve High, Phil Woodman, Martin Lowe and Viki Allan for their warm support and laboratory space for a part of this study, Martin Lowe for the gift of *E.coli* BL21(DE3)-RIL [pRK753], Keith Gull for the gift of hybridoma cell line expressing Tat1 antibody, and Jennifer Winstanley for technical assistance in the early stages of the project. This work was supported by Cancer Research UK (Grant numbers C5759/A20971 and C5759/A27412).

## Appendix Figure legends

**Appendix Figure S1.**
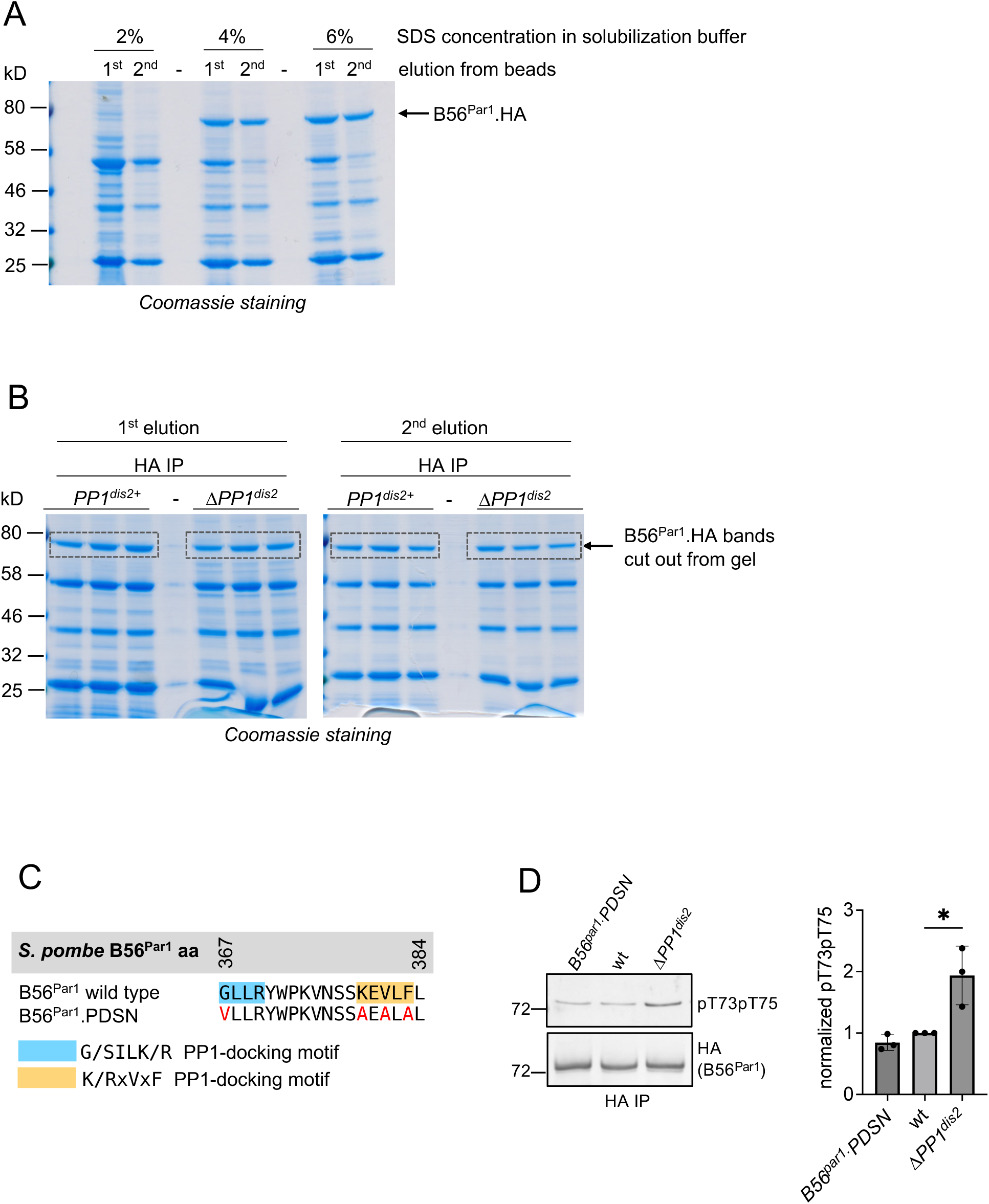
PP2A-B56^Par1^.T73T75 phosphorylation is PP1^Dis2^ dependent. (**A-B**) Affinity purification of B56^Par1^.HA for Quantitative Mass Spectrometry PTM analysis. (**A**) Optimization of HA-precipitation of the B56^Par1^ subunit under denaturing conditions (Grallert & Hagan, 2017a) from asynchronous B56^Par1^.HA *ura4.d18 leu1.32* cultures. Only buffers with 4-6% SDS concentrations could solubilize the precipitate. (**B**) B56^Par1^.HA precipitates from asynchronous cultures of *PP1^dis2+^* and *ΔPP1^dis2^* genetic backgrounds purified under optimized conditions (6% SDS in solubilization buffer) have been separated by SDS-PAGE and visualized by Coomassie staining. Boxes indicate B56^Par1^.HA bands that were excised for analysis. (**C**) Schematics of the PP1-docking site motifs in B56^Par1^ alongside B56^Par1^.PDSN PP1-docking site mutant (Grallert *et al*., 2015). (**D**) B56^Par1^.T73T75 phosphorylation levels are independent on PP1^Dis2^-docking site in B56^Par1^. T73T75 phosphorylation levels have been determined in B56^Par1^.HA-precipitates from indicated genetic backgrounds. Left, representative immunoblot; right, quantification of levels (means +/- SD, *n*=3). pT73pT75 levels were normalized to total levels of the B56^Par1^.HA target protein run on a parallel Western blot as sample processing controls, and normalized values were plotted relative to wild type, which was set to 1. Significance was calculated by a one-way ANOVA followed by Dunnett’s multiple comparisons test. *P < 0.05. wt, wild type.

**Appendix Figure S2.**
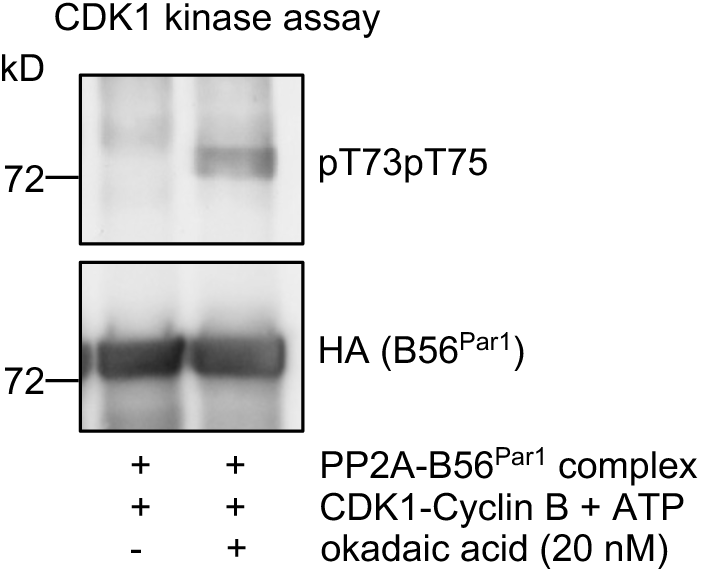
PP2A-B56^Par1^.T73T75 auto-dephosphorylates. PP2A-B56^Par1^.HA complexes were purified from asynchronous cultures and subjected to phosphorylation by CDK1-Cyclin B ± nM concentrations of okadaic acid to inhibit PP2A activity. Phosphorylation was detected by pT73pT75 phospho-specific antibodies with signal was developed by a chromogenic method, and the same membrane was then probed with anti-HA antibodies to detect B56^Par1^.HA as sample processing controls with signal developed by chemiluminescence.

**Appendix Figure S3.**
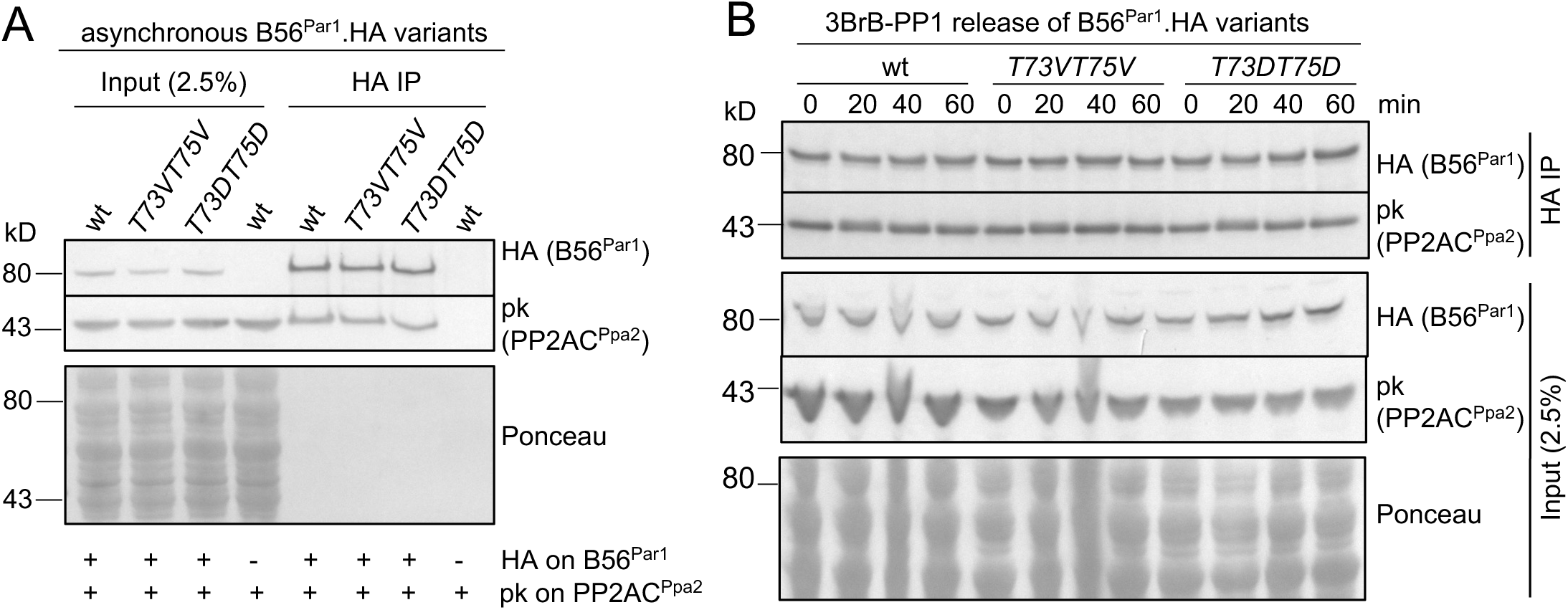
PP2A-B56^Par1^.T73T75 phospho-mutants maintain complex integrity. (**A-C**) B56^Par1^.T73XT75X.HA variants were HA-precipitated from asynchronous (**A**) or ATP analogue-synchronized (**B**) *ΔPP2Ac^Ppa1^* cultures and co-precipitation of the catalytic pk.PP2Ac^Ppa2^ subunit was monitored by Western blot. Ponceau staining was used as a loading control.

**Appendix Figure S4.**
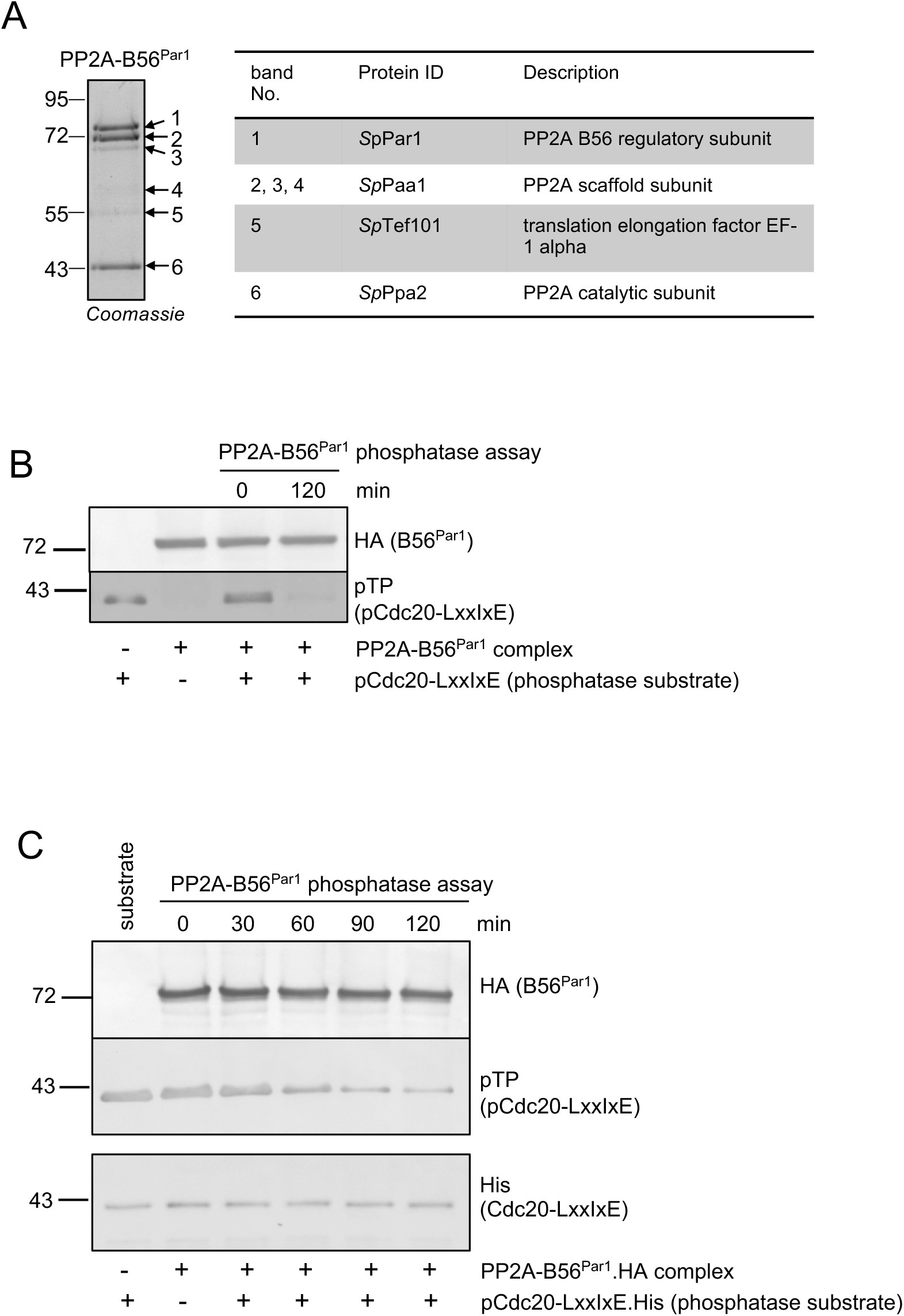
Endogenous asynchronous PP2A-B56^Par1^ complex purification and activity assay. (**A**) Purification of an endogenous PP2A-B56^Par1^.HA complex harboring the major pk.PP2AC^Ppa2^ subunit from asynchronous cultures. Left, Coomassie-stained protein gel. Right, protein identification (ID; most abundant exclusive unique spectrum count) of individual bands. (**B-C**) Activity assay of the endogenous PP2A-B56^Par1^ complex purified from asynchronous cells. Purified complexes were mixed with phosphorylated Cdc20-LxxIxE substrate and substrate dephosphorylation was monitored for 120 min alongside loading controls (B56^Par1^.HA) with samples withdrawn at intervals. A membrane probed with anti-His antibodies as sample processing control to detect total Cdc20.LxxIxE was processed in a parallel Western blot.

**Appendix Figure S5.**
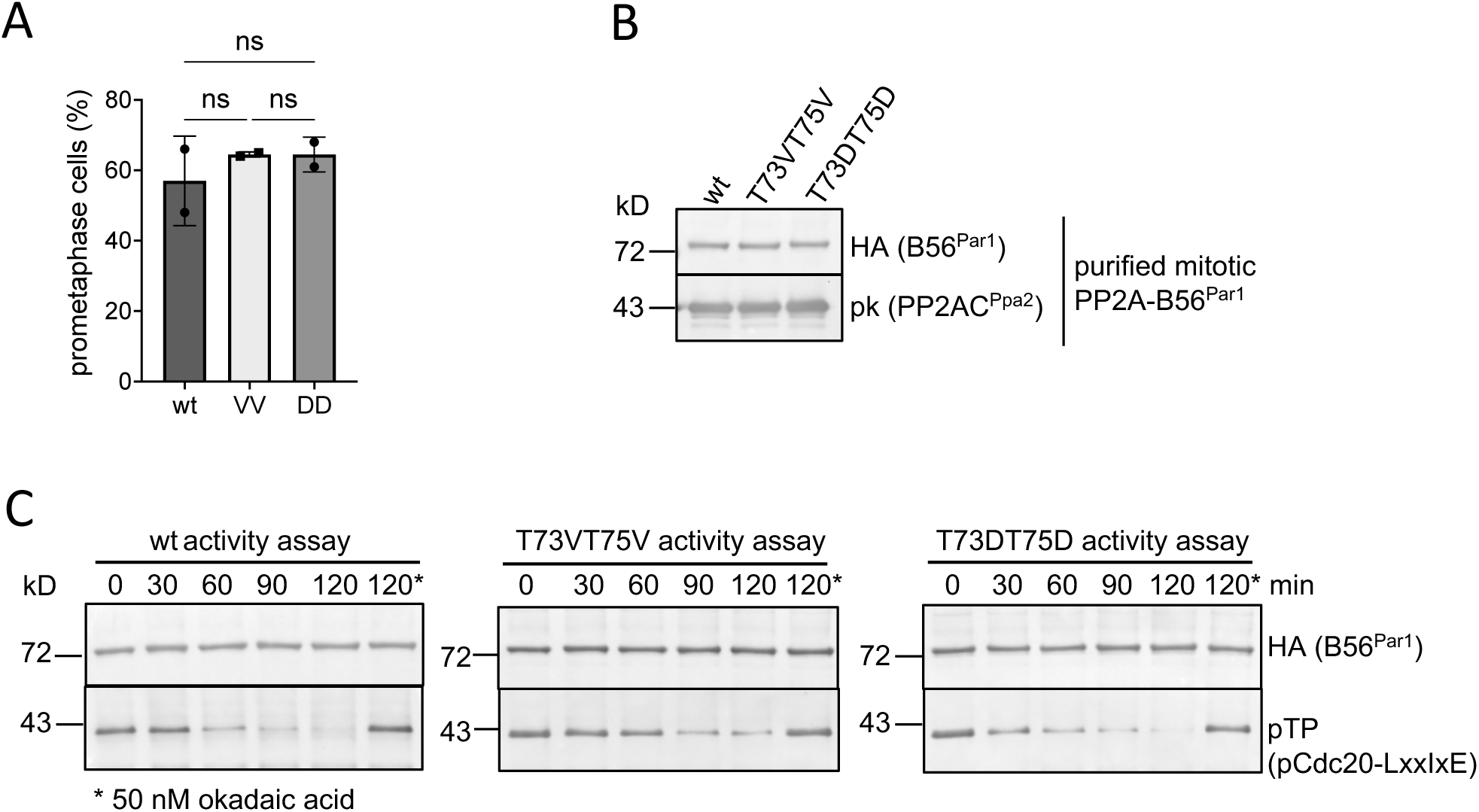
Endogenous mitotic PP2A-B56^Par1^.T73XT75X complexes activity assay. (**A**) *nda3.KM311*-induced mitotic arrest of *PP2A-B56^par1^.T73XT75X*. Non-significance was calculated by an ordinary one-way ANOVA followed by Turkey’s multiple comparisons test. (means +/- SD, *n*=2). (**B-C**) PP2A-B56^Par1^.T73T75 variants activity assay. PP2A-B56^Par1^ complexes were purified from mitotically arrested *nda3.KM311* cultures (**B**), mixed with phosphorylated Cdc20-LxxIxE substrate and substrate dephosphorylation was monitored alongside loading controls (B56^Par1^.HA) for 120 min with aliquots withdrawn at intervals (**C**). Phosphorylation was detected by pT73pT75 phospho-specific antibodies. * Okadaic acid (nM) treatment confirms a PP2A-specific dephosphorylation. T73VT75V, B56^Par1^.T73VT75V.HA complex; T73DT75D, B56^Par1^.T73DT75D.HA complex; wt, wild-type complex.

**Appendix Figure S6.**
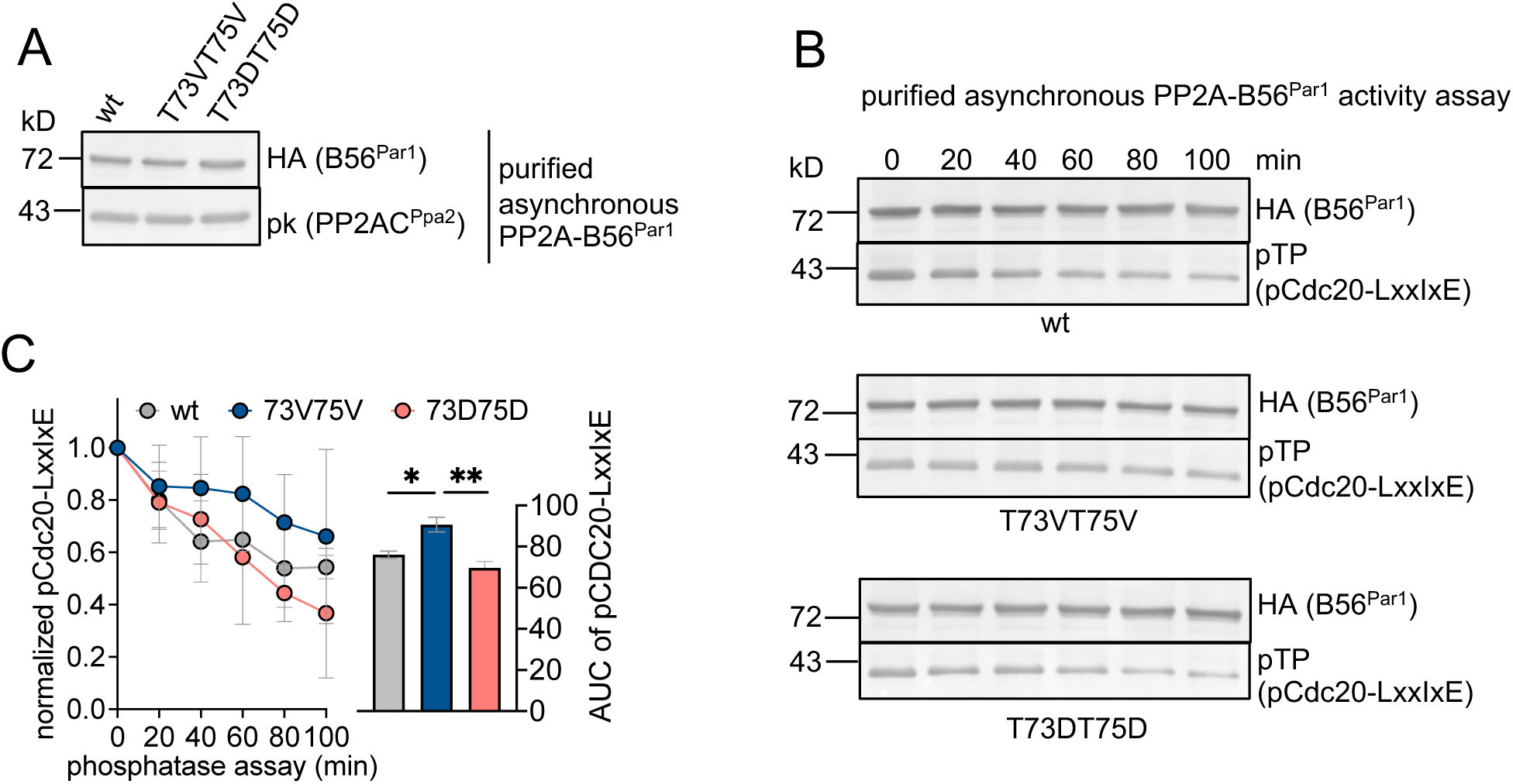
Endogenous asynchronous PP2A-B56^Par1^.T73XT75X complexes activity assay. (**A-B**) PP2A-B56^Par1^.73X75X.HA complexes variants purified from asynchronous cells (**A**) were mixed with phosphorylated Cdc20-LxxIxE substrate and substrate dephosphorylation was monitored for 100 min with samples withdrawn every 20 min (**B**). (**C**) Quantification of activity measured by level of substrate dephosphorylation (means +/- SD, *n*=3). pCdc20-LxxIxE levels were normalized against loading control (B56^Par1^.HA), and normalized values were plotted relative to time point 0 min, which was set to 1. Area under the curve (AUC) significance was calculated by unpaired *t*-test, *P < 0.05, **P < 0.01. T73VT75V, B56^Par1^.T73VT75V.HA complex; T73DT75D, B56^Par1^.T73DT75D.HA complex; wt, wild-type.

